# Development of a tightly regulated copper-inducible transient gene expression system in *Nicotiana benthamiana* incorporating suicide exon and Cre recombinase

**DOI:** 10.1101/2024.03.23.586378

**Authors:** Bing-Jen Chiang, Kuan-Yu Lin, Yi-Feng Chen, Ching-Yi Huang, Foong-Jing Goh, Lo-Ting Huang, Li-Hung Chen, Chih-Hang Wu

**Author notes:** These authors contributed equally to this work (Bing-Jen Chiang, Kuan-Yu Lin).

## Abstract

Chemical-inducible gene expression systems have been frequently used to regulate gene expression for functional genomics in various plant species. However, a convenient chemical-inducible system that can tightly regulate transgene expression in *Nicotiana benthamiana* is still missing. In this study, we developed a tightly regulated copper-inducible system that can be used to regulate transgene expression and perform cell death assays in *N. benthamiana*. We tested several chemical-inducible systems using *Agrobacterium*-mediated transient expression and found that the copper-inducible system showed the least concerns of leakiness issues. Using the MoClo-based synthetic biology approach, we optimized the design of the copper-inducible system and incorporated the use of the suicide exon HyP5SM/OsL5 and Cre/LoxP as additional regulatory elements to enhance the tightness of the regulation. This new design allowed us to tightly control the hypersensitive cell death induced by several tested NLRs and their matching AVRs, and it can also be easily applied to regulate the expression of other transgenes in transient expression assays. Our findings provide new approaches for both fundamental and translational studies in plant functional genomics.

## Introduction

Exploring gene function in plants relies on the use of various genetic tools to facilitate or disrupt the expression of the gene of interest. To understand the gene function or signaling events with a temporal resolution, several chemical-inducible systems have been developed to control the expression of transgenes (Zuo & Chua, 2000; Padidam, 2003; Moore *et al*., 2006; Borghi, 2010). Chemical-inducible systems typically involve chimeric transcription factors and corresponding promoters derived from genetic elements of heterologous organisms. This design prevents interference with the expression of endogenous genes and has been extremely useful in characterizing the function of different genes in various plant species (Padidam, 2003).

Several chemical-inducible systems have been developed and applied to plant biology research in the last few decades (Padidam, 2003). The dexamethasone (Dex)-inducible system uses a modified *Escherichia coli* lac-repressor mechanism with the chimeric transcription factor LhGR (Padidam, 2003). LhGR combines the lacIHis17 DNA binding domain, Gal4 transcription-activation-domain-II, and the DEX-binding domain of the rat glucocorticoid receptor (GR). The pOp6 promoter has a repeated binding site with six direct repeats of the lac operator (lacO) sequence. The introduction of DEX causes LhGR to move from the cytoplasm to the nucleus, activating the transcription of the downstream gene. The β-estradiol (β-EST)-inducible system is adapted from the *E. coli* LexA repressor system (Zuo *et al*., 2000). The synthetic transcription factor XVE is composed of bacterial repressor LexA (X), the acidic transactivating domain of VP16 (V), and the regulatory region of the human estrogen receptor (E). Upon β-EST application, XVE binds to the LexA operator to activate transcription of the gene fused to the operator. The tetracycline-(Tet-) inducible system is based on the Tet repressor (TetR) and the tet operator (tetO) elements of *E. coli* and has been modified into Tet-ON and Tet-OFF systems for turning gene expression ON or OFF, respectively (Padidam, 2003). In the Tet-ON system, the application of tetracycline or doxycycline (Dox) triggers a conformational change of TetR, leading it to bind to the tetO to activate downstream gene expression. The copper-inducible system originated from the copper-metallothionein regulatory system of *Saccharomyces cerevisiae* (Buchman *et al*., 1989; Mett *et al*., 1993; Saijo & Nagasawa, 2014). The copper-binding transcription factor CUP2 is fused to the viral transcription factor VP16. In the presence of copper, CUP2 undergoes conformational changes, enabling it to bind the operator sequence known as the Copper Binding Site (CBS) to activate the transcription of downstream genes. In addition to the systems mentioned above, several other methods, such as ethanol-, herbicide- and insecticide-inducible systems have also been developed to control transgene expression in plants (Padidam, 2003).

Plants utilize various immune receptors to detect pathogen molecules. In brief, plant cell surface-localized pattern recognition receptors (PRR) detect pathogen-associated molecular patterns (PAMPs) and apoplastic effectors, whereas intracellular nucleotide-binding domain leucine-rich repeat-containing proteins (NLRs) recognize cytoplasmic effectors secreted from pathogens (Ngou *et al*., 2022). The signaling events post pathogen recognition have been a long-standing interest in the plant biology research community. The utilization of chemical-inducible systems has provided valuable information for understanding the intricate signaling events post-immune activation. Several avirulence factors from *P. syringae*, including AvrRpt2, AvrPphB, AvrB and AvrRpm1, have been placed under Dex- or β-EST-inducible system for generating transgenic plants to understand immune responses or effector functions (McNellis *et al*., 1998; Nimchuk *et al*., 2000; Tornero *et al*., 2002; Dowen *et al*., 2009; Yuan *et al*., 2021). Recently, the “Super ETI” (SETI) Arabidopsis lines, which place *P. syringae* effector AvrRPS4 under LexA promoter, enable the activation of RPS4/RRS1-mediated immune responses after β-EST treatment (Ngou *et al*., 2020). The applications of chemical-inducible systems have allowed the dissection of NLR-mediated immunity without the activation of cell surface immune receptors, providing an in-depth understanding of NLR-mediated immunity and their cross-talk with PRR-mediated immunity (Ngou *et al*., 2021; Yuan *et al*., 2021).

While chemical-inducible systems have proven successful in generating transgenic lines for various plant research purposes, some studies have reported issues with leakiness (Okuzaki *et al*., 2011; Park *et al*., 2012; Caddell *et al*., 2015; Gonzalez *et al*., 2015; Garcia-Perez *et al*., 2022). Since the detection of immune receptors to corresponding AVRs is often highly sensitive, the leakiness issue can become particularly problematic in cases where NLR-type plant immune receptors are co-expressed with AVRs. For example, Gonzales et. al. (2015) mentioned that attempts to generate Bs2/AvrBs2 *N. benthamiana* and RPP1/ATR1 *A. thaliana* using the DEX-inducible system were unsuccessful (Gonzalez *et al*., 2015). Furthermore, in Gantner et al. (2018) where AvrRPS4 was placed under a UAS promoter and transferred into *N. benthamiana* through agroinfiltration, the protein accumulation of AvrRPS4 was detectable without DEX treatment, indicating the sign of leakiness before inducer application (Gantner et al., 2018).

To tackle this issue, Gonzales et al. (2015) employed the use of the suicide exon HyP5SM, a hybrid plant 5S rRNA mimic element derived from rice and Arabidopsis, in the target gene. In the absence of the rice splicing factor OsL5, the splicing variant introduces a premature stop codon, triggering nonsense-mediated decay (Hickey *et al*., 2012). When OsL5 is present, alternative splicing skips the suicide exon, producing the desired transcripts for gene expression (Hickey *et al*., 2012). The integration of HyP5SM suicide exon and OsL5 with the DEX-inducible system was successfully applied to genes such as GFP, AvrBs2, and ATR1 to tightly regulate protein expression (Hickey *et al*., 2012; Gonzalez *et al*., 2015). In experiments where both OsL5 and AvrBs2-HyP5SM or ATR1-HyP5SM were placed under the 6xUAS promoter and transiently transformed into *Nicotiana* spp. alongside constitutively expressed Bs2 or RPP1, cell death phenotypes were observed only after DEX treatment (Gonzalez *et al*., 2015). However, the limitation of this system is its dependency on inserting HyP5SM into the gene of interest at positions encoding Glu, Gln, or Lys, necessitating a case-by-case testing of potential insertion sites which can be a laborious process (Gonzalez *et al*., 2015).

While exploring the potential chemical-inducible systems for controlling cell death induced by NLR and AVR in plants, the copper-inducible system caught our attention. This system was successfully applied to control the expression of dCas9:EDLL and MS2:VPR for transcriptional activation of genes in *N. benthamiana* recently (Garcia-Perez *et al*., 2022). Here, to develop a tightly regulated chemical-inducible system for fine-tuning transgene expression in *N. benthamiana*, we evaluated various chemical-inducible systems for driving the expression of RUBY and luciferase (Contag & Bachmann, 2002; He *et al*., 2020), as well as conducting cell death assays in *N. benthamiana* through agroinfiltration. Our results showed that the copper-inducible system exhibited the least concerns of leakiness compared to other systems tested, and protein expression can be efficiently induced upon copper infiltration. To enhance the level of induction, we utilized MoClo-compatible modules and identified a combination that demonstrated higher reporter expression compared to the original design. To improve the tightness of the inducible system, we introduced the fluorescence protein mCherry2-HyP5SM and LoxP-mCherry-LoxP as MoClo N-terminal modules, along with the corresponding inducible splicing factor OsL5 and recombinase Cre. The incorporation of these additional regulatory elements allowed us to control very sensitive cell death mediated by NLRs and AVRs. This novel design has the potential for broad application in functional studies of plant genes using agroinfiltration-based experimental designs on *N. benthamiana* as well as in other experimental systems.

## Materials and Methods

### Plant growth condition

Wild type (WT) *N. benthamiana* were propagated in a walk-in chamber with a temperature 24–26°C, humidity of 45–65%, and 16/8 hr light/dark cycle.

### Agroinfiltration and chemical induction

Plasmids were transformed into *Agrobacterium tumefaciens* GV3101 using electroporation, and then spread on agar plate of 523 medium (1% sucrose 0.8% Casein hydrolysate (N-Z-Case) 0.4% Yeast extract 0.2% K_2_HPO_4_ and 0.03% MgSO_4_ 7H_2_O pH6.9) containing appropriate antibiotics. Two days later, colony PCR was performed to confirm the presence of the plasmid in the strain. Agroinfiltration experiments were performed using 4-week-old *N. benthamiana*. Strains of *A. tumefacien*s were refreshed from glycerol stock to 523 medium containing appropriate antibiotics at 28 °C overnight. Cells were harvested by centrifugation at 2500 × g, room temperature for 5 min. Subsequently, cells were resuspended in MMA buffer (10 mM MgCl2, 10 mM MES-KOH, 150 µM acetosyringone, pH5.6) to the OD_600_ of 0.2 and then infiltrated into leaves using 1 mL syringes. For chemical induction, solutions of 10μM CuSO_4_ (50μM β-estradiol, 2.25μM doxycycline, 30μM dexamethasone, or as indicated for testing suitable inducible system) were infiltrated on the agroinfiltrated sites using 1 ml syringes at 1 day-post-infiltration of *A. tumefaciens*.

### RUBY quantification

Leaf discs were sampled by using a cork borer with a 0.5 cm diameter at 36 hrs after chemical induction. Each leaf disc was then mounted with 200 μL extraction buffer (10% ethanol and 0.1% formic acid) in a 96-well plate. After 2 hours of gentle shaking, 150 μL of the extraction buffer was transferred to a new 96-well plate, and the absorbance was measured using a spectrophotometer (Agilent BioTek Synergy H1) at wavelengths of 475, 535 and 600 nm. The RUBY absorbance was calculated by using the following formula: (OD_475_ - OD_600_) + (OD_535_ - OD_600_) (Polturak *et al*., 2017).

### Luciferase assay

At 36 hours after chemical induction, leaf discs were sampled using a cork borer with a 0.5 cm diameter and immersed in 200 μL distilled water in a white 96-well plate. After removing the distilled water, 200 μL of 471 μM D-luciferin (LUCK-100, GoldBio) was added to each well and incubated for 10 minutes in the dark. The relative luminescence units (RLU) were detected using the IVIS Lumina LT In Vivo Imaging System (Revvity) with a 2-second exposure time.

### Cell death quantification

NLRs and/or corresponding effectors were transiently expressed using the copper-inducible system via *A. tumefaciens*-mediated protein expression. At 36 hours post-copper treatment, the intensity of cell death was quantified using the UVP ChemStudio Imaging Systems (Analytik Jena) as described previously (Goh *et al*., 2023). In brief, raw autofluorescence images of cell death were acquired using blue LED light for excitation and a FITC filter (513 - 557 nm) as the emission filter. The mean signal intensity of the selected infiltrated area was further normalized with the maximum intensity (65535) to obtain the relative intensity of cell death.

### Plasmid construction and gene synthesis

All constructs used in this study were generated using Golden Gate assembly following the MoClo system (Engler *et al*., 2008, 2014; Weber *et al*., 2011; Gantner *et al*., 2018). Level 0 modules of TetO, CBS4-miniDFR, TetRv10, CUP2, AVR3a_KI, loxP-mCherry, VP16, VP64, p65, and GAL4-AD were synthesized as MoClo-compatible units in kanamycin-resistant pUC57 vectors using the services provided by SynBio Technologies (New Jersey, USA). Sequences of plasmids, modules of the inducible system, NLRs, AVRs, and primers used in this study are listed in Table S1-2.

### Protein extraction and western blot analysis

SDS-PAGE electrophoresis and western blot analyses were conducted as previously described with minor modifications (Win *et al*., 2011). Briefly, 16 hours after copper treatment, leaf discs were collected into 2 ml microtubes containing ceramic beads and flash-frozen with liquid nitrogen. Subsequently, the leaf tissues were disrupted using the SH-100 homogenizer (J&H Technology Co., Ltd.) at 800 rpm for 30 seconds, repeated five times. Total protein extraction was then performed using extraction buffer composed of 10% glycerol, 25 mM Tris pH 7.5, 1 mM EDTA, 150 mM NaCl, 2% w/v PVPP, 10 mM DTT, 1x protease inhibitor cocktail (Sigma), and 0.1% IGEPAL (Sigma). After centrifugation at 13,000 x g for 10 minutes at 4°C, the supernatant fraction was collected and mixed with 4x sample loading dye (200 mM Tris-HCl pH 6.8, 8% (w/v) SDS, 40% (v/v) glycerol, 50 mM EDTA, 0.08% bromophenol blue and 100 mM DTT). The protein samples were then incubated at 70°C for 10 minutes prior to SDS-PAGE electrophoresis and immunodetection analyses. Primary antibodies used included anti-myc (A00704, GenScript), anti-RFP (YH80520, Yao-Hong), or anti-flag (F1804, Sigma), as indicated, with Peroxidase-conjugated AffiniPure Goat Anti-Mouse IgG (H+L) (115-035-003, Jackson) or Peroxidase Conjugated Goat Anti-Rabbit IgG (H+L) (AP132P, Sigma) as secondary antibodies. Images were captured using the UVP ChemStudio Imaging Systems (Analytik Jena). SimplyBlue SafeStain (465034, Invitrogen) was used to visualize the amount of rubisco on the PVDF membrane.

## Results

### The copper-inducible system enables effective induction of gene expression in *N. benthamiana*

To identify a chemical-inducible gene expression system suitable for transient expression in *N. benthamiana*, we evaluated four previously reported methods, including the β-EST-, DEX-, Tet-, and copper-inducible systems, in the regulation of reporter gene expression. We fused the promoters (LexA, pOp6, tetO, or CBS4) to the RUBY reporter gene or firefly luciferase. One day after the *Agrobacterium*-mediated co-infiltration of the transcription factor (XVE/LhGR/TetR/CUP2) together with the matching promoter driving RUBY or luciferase, we treated the plants with the corresponding chemicals (β-EST/DEX/Dox/Copper) using syringe infiltration at the previous infiltration sites. We collected the leaf discs for RUBY quantification at 36 hours after chemical treatment, or for luciferase assays at 24 hours after chemical treatment. Although DEX- and β-EST-inducible systems show high reporter expression after chemical treatments, both systems showed very strong leakiness issues without chemical treatment (Figure 1A and 1B). While the Tet-inducible system showed a low level of leakiness, the efficiency of the induction was not as good as other systems. The copper-inducible system showed no sign of leakiness and can be induced by the treatment of copper sulfate (Figure 1A and 1B). To clarify whether this leakiness issue is due to the inducible promoter itself or the inducible promoter together with the matching transcription factor, we infiltrated the reporter genes driven by the inducible promoter with and without the matching transcription factors in the absence of the corresponding chemicals. We found that co-infiltration of inducible promoters together with the matching transcription factor led to the leakiness issue in the β-EST- and DEX-inducible systems, suggesting that the leakiness issue is due to the overexpression of the transcription factor in *N. benthamiana*. However, this issue was not observed in the Tet- and copper-inducible systems (Figure S1).

**Figure 1.**
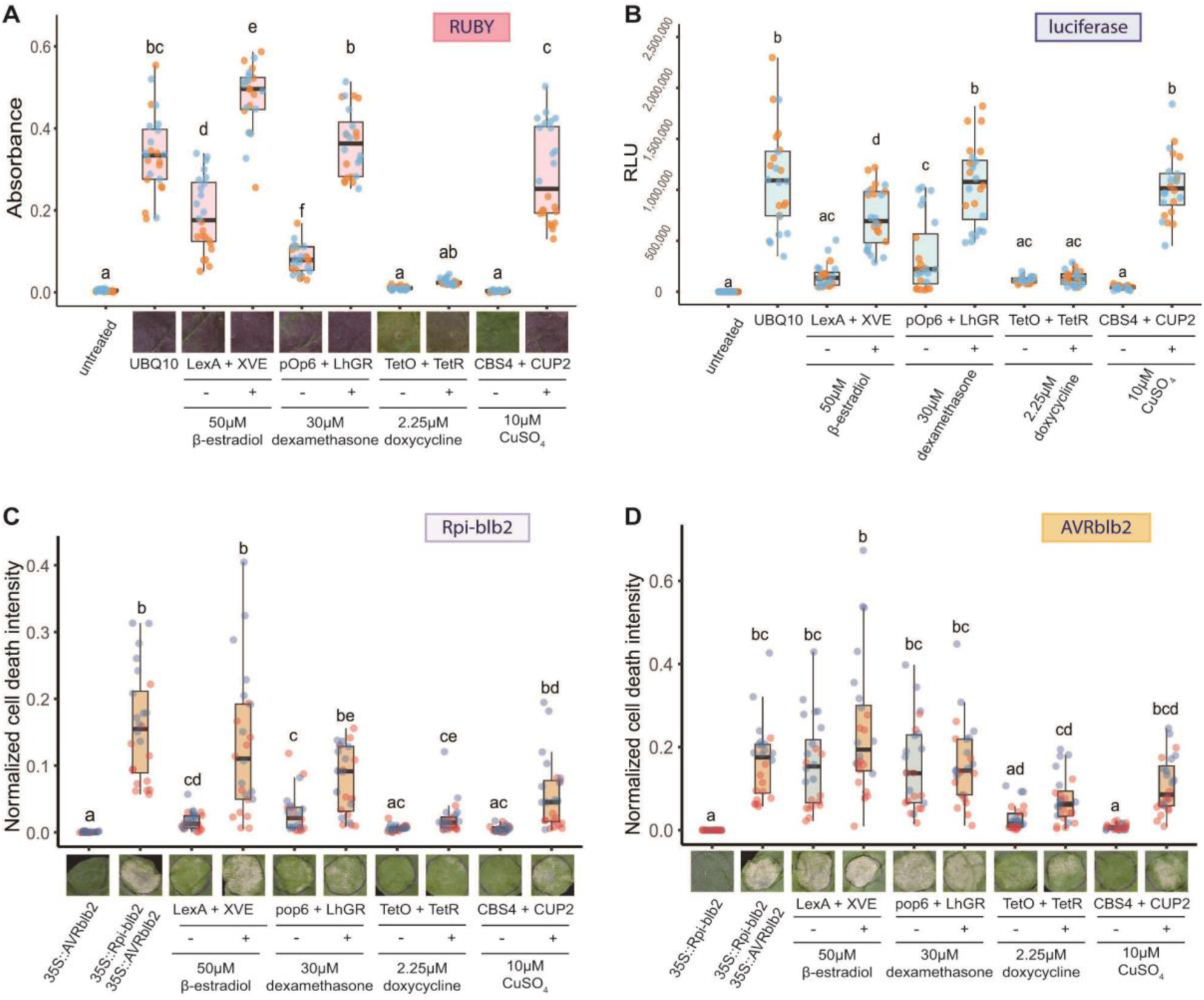
The copper-inducible system stands out as an optimal choice for regulating gene expression in *Nicotiana benthamiana*. Measurement of RUBY absorbance (A) and luciferase activity (B) controlled by the β-estradiol, dexamethasone, doxycycline, or copper-inducible systems in *N. benthamiana*. RUBY and luciferase reporters were placed under the control of the UBQ10 promoter (as a positive control) or the respective inducible promoters and co-expressed with the corresponding transcription factor. After 24 hours, the corresponding chemicals were infiltrated to stimulate gene expression. For the RUBY assays, photographs were taken at 36 hours post-chemical induction, followed by RUBY absorbance measurement. Luciferase assays were performed at 24 hours post-chemical induction. Cell death triggered by the inducible Rpi-blb2 (C) or inducible AVRblb2 (D) in *N. benthamiana*. Chemicals were infiltrated at 24 hours post-agroinfiltration. Cell death intensity was quantified using UVP ChemStudio at 36 hours after chemical infiltration. Dots with different colors represent the results from independent biological replicates. Statistical analysis was conducted using TUKEY’s HSD test (p < 0.05) for the RUBY and luciferase assays, and Dunn’s test (p < 0.05) for the cell death assays.

To test whether these chemical-inducible gene expression systems can be used to control cell death induced by the recognition of pathogen effectors through intracellular immune receptors, we fused the inducible promoters to Rpi-blb2, a solanaceous NLR-type immune receptor, or AVRblb2, a *Phytophthora infestans* effector recognized by Rpi-blb2. One day after the co-infiltration of these constructs, we treated the plant leaves with corresponding chemicals. We quantified the cell death intensity at 36 hours after chemical treatment using autofluorescence-based imaging. Similar to the previous results, the copper-inducible system was the only system that showed no leakiness issue and can be successfully induced after chemical treatment (Figure 1C for inducible Rpi-blb2; and Figure 1D for inducible AVRblb2). Based on the results above, we decided to further optimize the copper-inducible system for *N. benthamiana*.

### Optimization of the copper-inducible system in *N. benthamiana*

To further optimize the copper-inducible system in *N. benthamiana*, we first tested the concentration of copper sulfate sufficient to trigger the reporter gene expression. We used the RUBY reporter for optimization as it is a visual reporter that can be easily quantified. We tested 7 different concentrations of copper sulfate, ranging from 0.1 to 50µM, in inducing RUBY expression. While treatments with low concentrations of copper sulfate, such as 0.5µM and 1µM, were sufficient to induce reporter gene expression, the RUBY signals were usually very patchy with stronger phenotypes near the copper-infiltrated sites (Figure 2A). Treatments with copper sulfate higher than 10µM showed no clear differences in RUBY expression. However, we occasionally observed necrotic symptoms with 25µM or 50µM copper sulfate treatments (Figure 2B). Therefore, we used 10µM copper sulfate as a standard treatment for the follow-up experiments.

**Figure 2.**
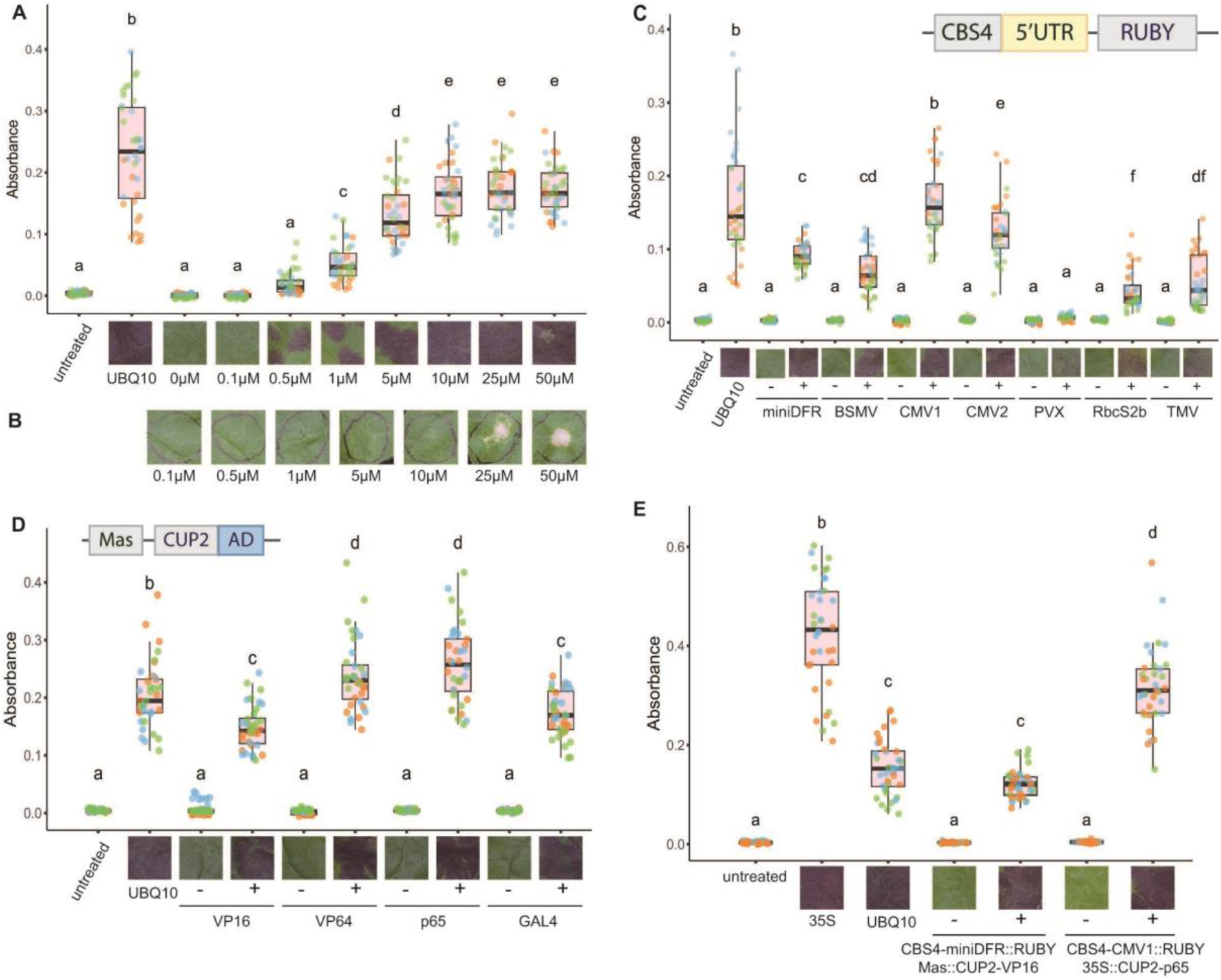
Optimization of the copper-inducible system. **(A)** Identification of the optimal concentration of CuSO_4_ for the copper-inducible system. *A. tumefaciens* strains harboring CBS4-miniDFR::RUBY and Mas::CUP2-VP16 were co-infiltrated in four-week-old *N. benthamiana*. The plants were then syringe-infiltrated with different concentrations of CuSO_4_ at 24 hours post agroinfiltration. **(B)** Photographs of the necrosis triggered by CuSO_4_ infiltration. Different concentrations of copper sulfate were infiltrated into the leaves of 4-week-old *N. benthamiana*. **(C)** Comparisons of different 5’UTR connecting to the CBS4 promoter in the copper-inducible system. *A. tumefaciens* strains harboring CBS4 promoter connecting to different 5’UTR and RUBY, and Mas::CUP2-VP16 were co-infiltrated in four-week-old *N. benthamiana*. **(D)** Comparisons of different activation domains fused to CUP2 in the copper-inducible system. *A. tumefaciens* strains harboring CBS4-miniDFR::RUBY and Mas::CUP2 fused to different activation domains were co-infiltrated in four-week-old *N. benthamiana*. **(E)** The combination of CMV1/35S/p65 modules induced a higher level of RUBY expression compared to the combination of miniDFR/Mas/VP16 modules. UBQ10::RUBY or 35S::RUBY were used as positive controls. The plants were then syringe-infiltrated with 10µM CuSO_4_ at 24 hours post agroinfiltration. Photographs were taken at 36 hours after CuSO_4_ infiltration, followed by RUBY absorbance measurement. Dots with different colors represent the results from independent biological replicates. Statistical differences were performed by Tukey’s HSD test (p<0.05).

Next, we tested the effect of 5’UTR (also referred to as minimum promoter) on the level of induction after copper treatment. In addition to the miniDFR that was used in the previous assays, we fused the CBS4 promoter to 6 different 5’UTR modules available in the MoCloPlant kit. We found that changing the 5’UTR to CMV1 or CMV2 increased the level of RUBY reporter expression after chemical treatment (Figure 2C). Then, we tested the effect of the promoters in driving the expression of CUP2. We changed the promoter from Mas to five other promoter modules and found that all these five other promoters performed better than the Mas promoter (Figure S2). We then tested whether changing the activation domain from VP16 to others can improve the efficiency of induction. We found that VP64 and p65 outperform VP16 in reporter gene induction when fusing to CUP2 (Figure 2D).

To test whether the elements identified above improve the expression of the RUBY reporter, we generated CBS4-CMV1::RUBY and 35S::CUP2-p65 constructs and compared them to the CBS4-miniDFR::RUBY and Mas::CUP2-VP16 that were used in our earlier experiments. We found that, indeed, the new combination (CBS4-CMV1::RUBY and 35S::CUP2-p65) showed significantly higher RUBY reporter expression compared to the original design (Figure 2E). Furthermore, this enhanced copper-inducible system consistently outperformed the level of expression driven by the constitutive promoter UBQ10 (Figure 2E).

### The copper-inducible system established above can be used to conduct some but not all cell death assays tested

To test whether the modified design identified above can be used to regulate cell death induced by NLR proteins and AVR proteins, we used the CBS4-CMV1 promoter to drive the expression of either Rpi-blb2 or AVRblb2 and compared that to the original condition tested in Figure 1. We found that the combination of CBS4-CMV1::Rpi-blb2/35S::AVRblb2/35S::CUP2-p65 induced occasionally stronger cell death than the original condition tested, although the statistical analysis revealed no significant differences due to the batch variation (Figure 3A). Similarly, the combination of CBS4-CMV1::AVRblb2/35S::Rpi-blb2/35S::CUP2-p65 occasionally induced stronger cell death than the original condition, although the statistical analysis was not significant (Figure 3B). These results indicated that the modified copper-inducible system can also be used to drive the expression of both Rpi-blb2 and AVRblb2 for cell death analysis.

**Figure 3.**
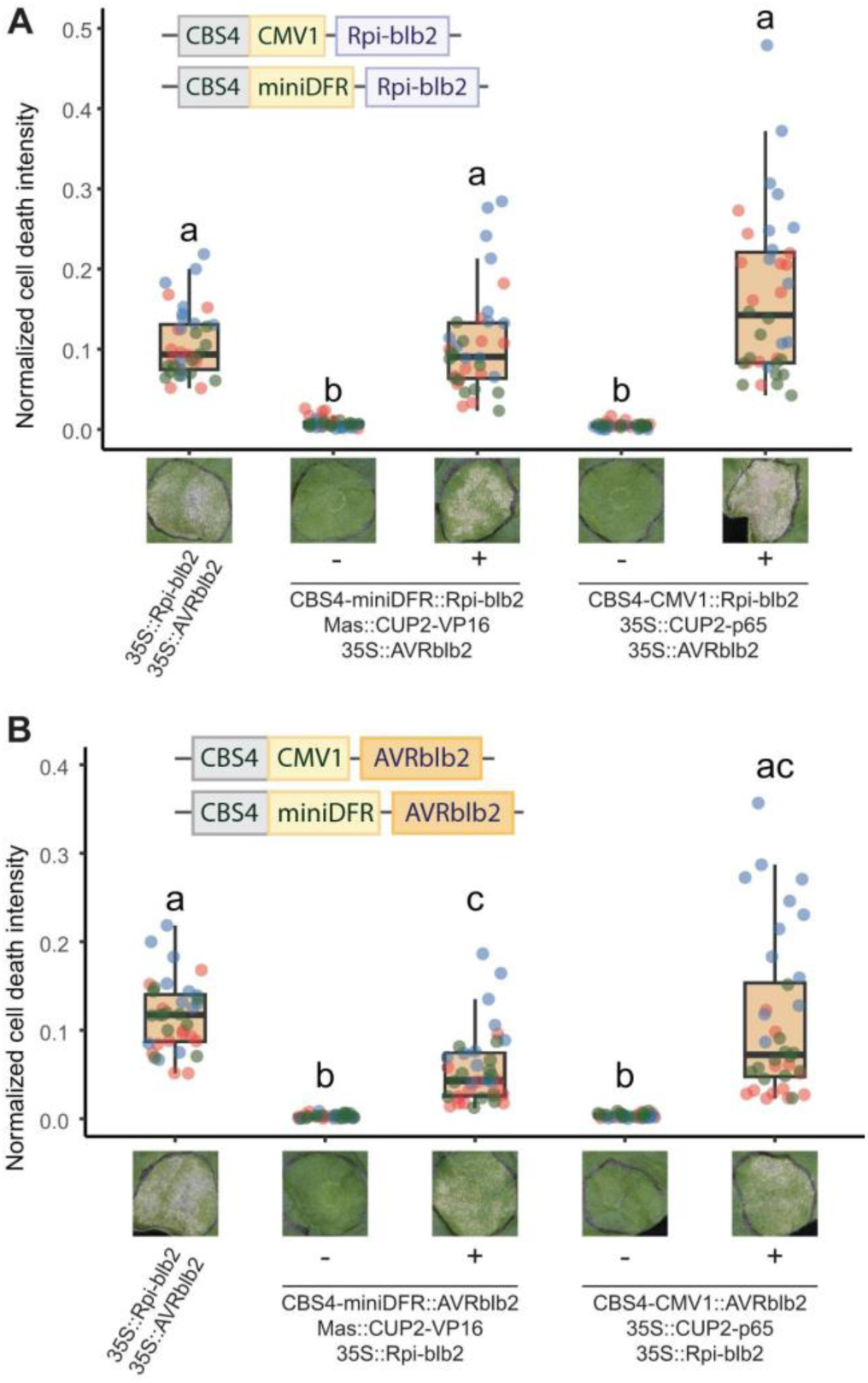
The copper-inducible system can regulate cell death triggered by Rpi-blb2 and AVRblb2. (A) Cell death assays comparing Rpi-blb2 driven by the constitutive 35S, CBS4-miniDFR or CBS4-CMV1 promoters, together with Mas::CUP2-VP16 or 35S::CUP2-p65 and 35S::AVRblb2. (B) Cell death assays comparing AVRblb2 driven by the constitutive 35S, CBS4-miniDFR or CBS4-CMV1 promoters, together with Mas::CUP2-VP16 or 35S::CUP2-p65 and 35S::Rpi-blb2. Dots with different colors represent the results from independent biological replicates. Statistical differences were performed by Dunn’s test (p<0.05).

To further test whether the copper-inducible system can regulate cell death induced by other NLRs and/or their corresponding AVRs, we used the inducible promoter to drive the expression of NRC4^DV^ (an autoactive variant of the helper NLR NRC4), and Rx/CP pair (Solanaceous NLR Rx and the matching AVR coat protein of *Potato Virus X* (PVX)), respectively. We found that the cell death induced by NRC4^DV^ was observed only after copper treatment (Figure 4A). However, clear cell death phenotypes were observed when we used the inducible system to drive the expression of Rx or CP (Figure 4B). We reasoned that this might be due to the detection of CP by the NLR protein Rx being extremely sensitive or the two proteins exhibiting slow turnover rates, and thus the experiment tolerant very low levels of leakiness. Further modifications are required to enable the application of the copper-inducible system to regulate cell death induced by Rx/CP.

**Figure 4.**
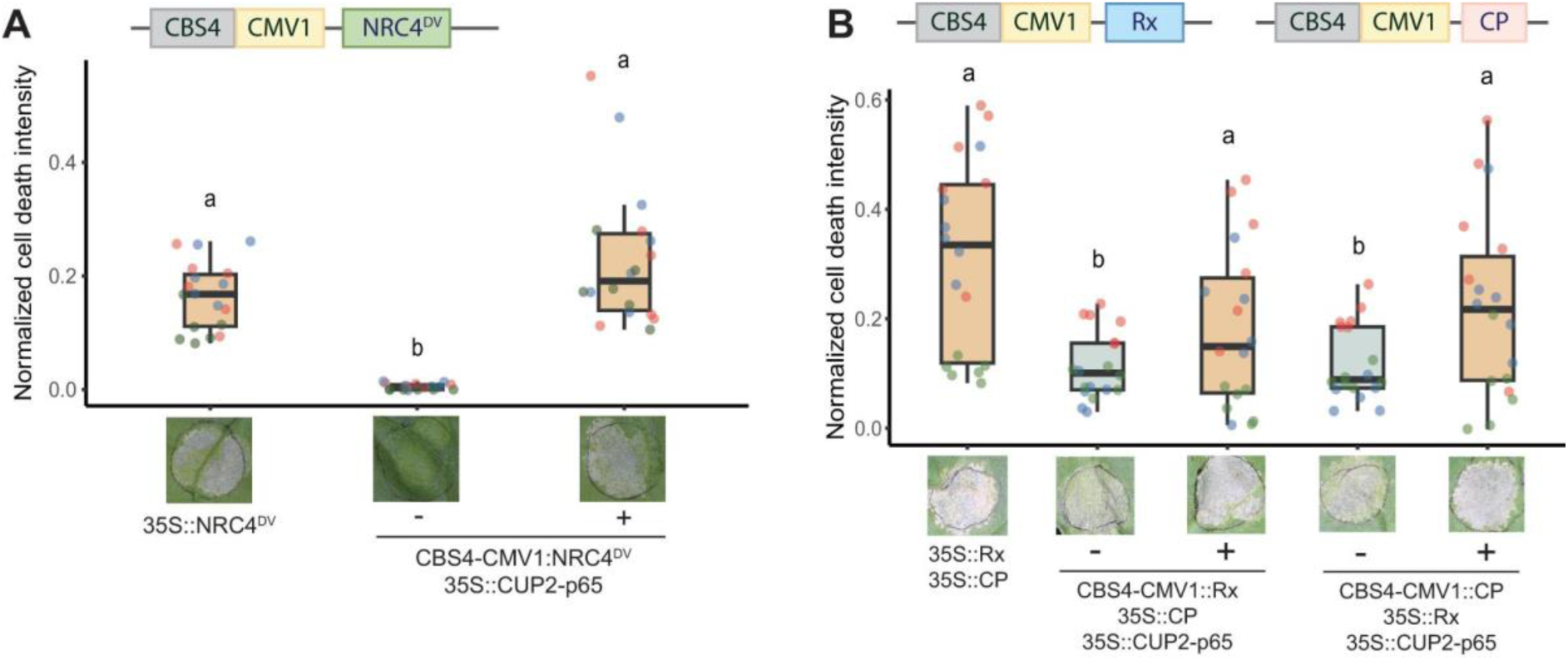
The copper-inducible system can regulate cell death caused by NRC4^DV^ but not Rx/CP. (A) Cell death assay of the copper-inducible expression of NRC4^DV^. 35S::NRC4^DV^ (as positive control) and CBS4-CMV1::NRC4^DV^/35S::CUP2-p65 were agroinfiltrated on *N. benthamiana* leaves. (B) Cell death assay of the copper-inducible expression of Rx and CP. 35S::Rx or 35S::CP (as positive control) and CBS4-CMV1::Rx or CBS4-CMV1::CP were agroinfiltrated with 35S::CUP2-p65 on *N. benthamiana* leaves. The plants were syringe-infiltrated with 10µM CuSO_4_ at 24 hours post agroinfiltration. Cell death intensity was quantified using UVP ChemStudio at 36 hours post-copper treatment. Dots with different colors represent the results from independent biological replicates. Statistical differences were performed by Dunn’s test (p<0.05).

### Incorporation of the alternative splicing cassette HyP5SM to improve the regulation of the copper-inducible system

To tackle this issue mentioned above, we incorporated the use of suicide exon HyP5SM which was previously applied to solve the leaky issue of Bs2/AvrBs2 and RPP1/ATR1 in the DEX-inducible system (Gonzalez *et al*., 2015). However, in the previous report, suicide exon HyP5SM was inserted directly into AvrBs2 or ATR1 genes, and this required tedious steps of testing the possible HyP5SM insertion sites (Gonzalez *et al*., 2015). Modified from the previous report inserting HyP5SM into the green fluorescence protein (Hickey *et al*., 2012), we inserted the HyP5SM in-between one of the Glutamic acid/Arginine sites in the red fluorescence proteins mCherry2 and then used this mCherry2-HyP5SM as an N-terminal tag module compatible with the MoClo system (Figure 5A). We then used the CBS4 promoter to drive the expression of both mCherry2-HyP5SM-CDS and OsL5, a factor essential for proper splicing of the HyP5SM suicide exon (Figure 5A).

**Figure 5.**
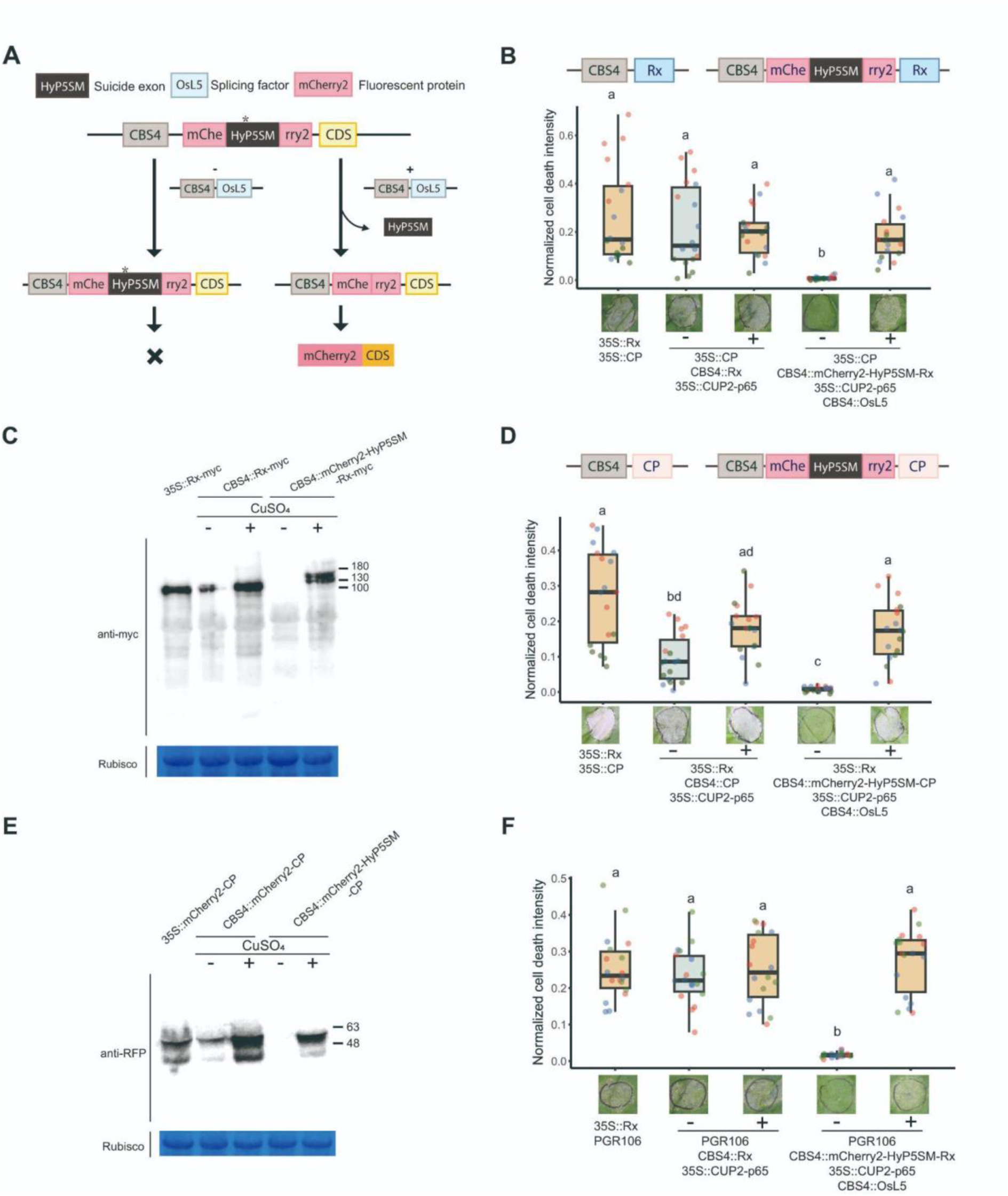
Incorporation of mCherry2-HyP5SM/OsL5 to improve the regulation of the copper-inducible system. (A) Design of copper-inducible mCherry2-HyP5SM and OsL5 system. HyP5SM is inserted into mCherry2 and designed as an N-terminal tag MoClo module. Both mCherry2-HyP5SM-CDS and OsL5 are driven by the CBS4 promoter. HyP5SM can be properly spliced by OsL5 only after copper treatment, leaving an mCherry2 tag fusing to the CDS. (B) Cell death assay of inducible mCherry2-HyP5SM-Rx/OsL5 with 35S::CP. *A. tumefaciens* strains harboring indicated expression constructs were co-infiltrated in four-week-old *N. benthamiana*. (C) Western blot of Rx-myc driven by 35S promoter, CBS4 promoter, and CBS4 promoter with mCherry2-HyP5SM/OsL5 regulation. Samples were collected at 24 hours post copper treatment. Proteins were detected using α-myc antibody, and SimplyBlue SafeStain-staining of Rubisco was used as the loading control. (D) Cell death assay of inducible mCherry2-HyP5SM-CP/OsL5 with 35S::Rx. *A. tumefaciens* strains harboring indicated expression constructs were co-infiltrated in four-week-old *N. benthamiana*. (E) Western blot of mCherry2-CP driven by 35S promoter, CBS4 promoter, and CBS4 promoter with mCherry2-HyP5SM/OsL5 regulation. Samples were collected at 24 hours post copper treatment. Proteins were detected using α-RFP antibody, and SimplyBlue SafeStain-staining of Rubisco was used as the loading control. (F) Cell death assay of inducible mCherry2-HyP5SM-Rx/OsL5 with PVX (pGR106). *A. tumefaciens* strains harboring indicated expression constructs were co-infiltrated in four-week-old *N. benthamiana*. The plants were syringe-infiltrated with 10µM CuSO_4_ at 24 hours post agroinfiltration. Cell death intensity was quantified using UVP ChemStudio at 36 hours post-copper treatment. For the cell death assays, dots with different colors represent the results from independent biological replicates. Statistical differences were performed by Dunn’s test (p<0.05).

To test whether this design can resolve the leakiness issue observed in inducible Rx experiments, we generated copper inducible mCherry2-HyP5SM-Rx and co-agroinfiltrated with inducible OsL5 and constitutively expressed CP. We found that the incorporation of mCherry2-HyP5SM significantly reduced the leakiness issue in the cell death assays with inducible Rx (Figure 5B). While western blot analysis revealed detectable Rx accumulation when driven by the CBS4 promoter, no Rx protein signal was detected when the regulation by mCherry2-HyP5M and inducible OsL5 was incorporated without copper treatment (Figure 5C). Next, we fused mCherry2-HyP5SM to CP and tested whether this construct showed any leakiness before copper treatment. Consistent with the previous observation, this design resolved the leakiness issue observed in the inducible CP experiments, with the cell death phenotype only observed after copper treatment (Figure 5D). Consistently, western blot analysis revealed detectable CP accumulation when driven by the CBS4 promoter, and this leakiness issue was fixed when the regulation by mCherry2-HyP5M and inducible OsL5 were incorporated (Figure 5E).

To further test whether this method can be combined with pathogen inoculation assays, we performed PVX (pGR106) inoculation experiments together with inducible Rx and mCherry2-HyP5SM-Rx. We found that while co-infiltration of CBS4::Rx-myc with PVX showed strong cell death without copper treatment, the co-infiltration of CBS4::mCherry2-HyP5SM-Rx with PVX showed cell death only after copper treatment (Figure 5F). These results indicate that the incorporation of the mCherry2-HyP5SM/OsL5 alternative splicing machinery improved the tightness of the inducible system, allowing it to be used for very sensitive cell death and resistance induced by Rx and CP.

### Incorporation of Cre/LoxP recombination system to improve the regulation of the copper-inducible system

We further tested whether mCherry2-HyP5SM can be incorporated to regulate the cell death induced by other NLRs and AVRs, such as R3a and AVR3a (Solanaceous NLR R3a and the matching AVR effector of *P. infestans*). However, the N-terminal fusion of the fluorescence protein compromised the function of R3a, and thus the incorporation of mCherry2-HyP5SM did not work for inducible R3a experiments (Figure S3A). Furthermore, a fusion of mCherry2-HyP5SM to AVR3a did not fully activate cell death in our experimental attempts (Figure S3B). Since incorporating mCherry2 as an N-terminal tag is not ideal for some experimental designs, we decided to explore other approaches to improve the copper-inducible system. Inspired by the studies incorporating Cre/LoxP system for improving the regulation of heat-shock promoters (Harrington *et al*., 2020; Tomoi *et al*., 2023), we tested whether the Cre/LoxP recombination system can also be used to improve the copper-inducible system.

We synthesized two LoxP-mCherry variants as MoClo N-terminal tag modules, one containing a mCherry CDS followed by a 35S terminator flanked by two LoxP sites, and the other one containing a mCherry CDS with premature stop codons followed by a 35S terminator flanked by two LoxP sites (Figure 6A). We then used the CBS4 promoter to drive the expression of both LoxP-mCherry(*)-LoxP-CDS and Cre recombinase, which mediates the recombination of the two LoxP sites, leading to the expression of the CDS under copper treatment (Figure 6A).

**Figure 6.**
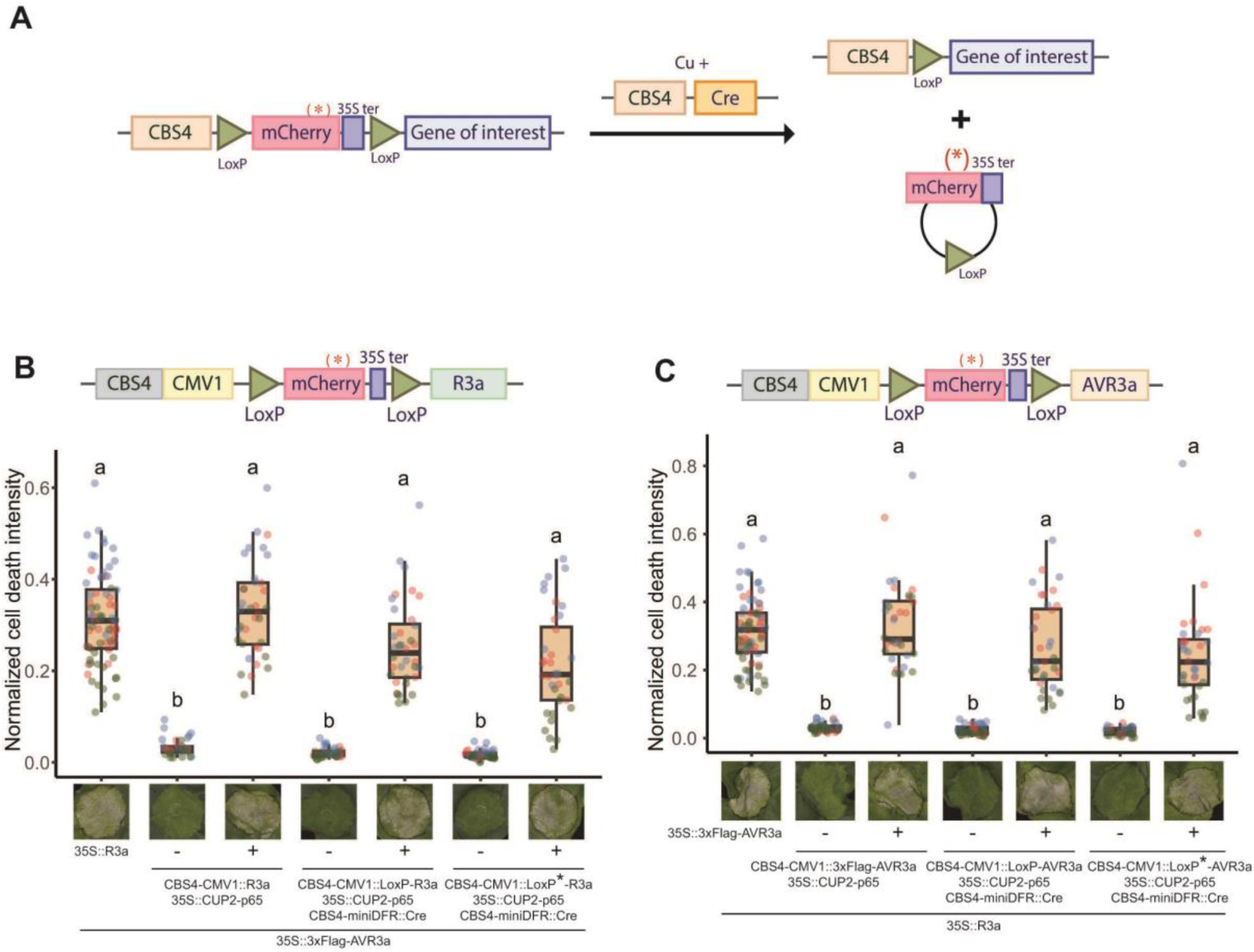
Incorporation of Cre/LoxP to improve the regulation of the copper-inducible system. (A) Design of copper-inducible Cre/LoxP system. mCherry-35Ster or mCherry*(premature stop codons)-35Ster was flanked by two LoxP sites and designed as an N-terminal tag MoClo module. Both LoxP-mCherry-LoxP-CDS and Cre are driven by the CBS4 promoter. Sequences between the two LoxP sites are removed upon copper treatment, leaving the expression of the CDS without interference from the mCherry tag. (B) Cell death assays of inducible LoxP-mCherry-LoxP-R3a/Cre with 35S::AVR3a. (C) Cell death assays of inducible LoxP-mCherry-LoxP-AVR3a/Cre with 35S::R3a. *A. tumefaciens* strains harboring indicated expression constructs were co-infiltrated in four-week-old *N. benthamiana*. The plants were syringe-infiltrated with 10µM CuSO_4_ at 24 hours post agroinfiltration. Cell death intensity was quantified using UVP ChemStudio at 36 hours post-copper treatment. For the cell death assays, dots with different colors represent the results from independent biological replicates. Statistical differences were performed by Dunn’s test (p<0.05).

To test whether this design can be used for inducible R3a experiments, we generated copper inducible LoxP-mCherry(*)-LoxP-R3a and co-agroinfiltrated with inducible Cre and constitutively expressed AVR3a. We observed clear cell death phenotype at 36 hours after copper treatment when either LoxP-mCherry-LoxP-R3a or LoxP-mCherry(*)-LoxP-R3a were used in the experiment. No clear cell death was observed without copper treatment (Figure 6B). Next, we fused LoxP-mCherry(*)-LoxP to AVR3a and tested whether cell death can be fully activated after copper treatment. For both designs, we observed a similar level of cell death intensity compared to using constructs with the 35S constitutive promoter or without Cre/LoxP regulation. No clear differences were observed in the absence of copper treatment (Figure 6C). These results suggest that the Cre/LoxP recombination system can be used as an alternative approach to the suicide exon to improve the regulation of copper-inducible system for cell death assays.

## Discussion

*N. benthamiana* has become a prominent experimental system in plant biology research (Bally *et al*., 2018; Ranawaka *et al*., 2023). Although numerous tools have been developed for assays in *N. benthamiana* (Bally *et al*., 2018; Derevnina *et al*., 2019), the absence of tightly regulated chemical-inducible systems has constrained certain experimental designs. In our study, we demonstrate the applicability of copper-inducible-based systems for controlling reporter gene expression and cell death assays through *Agrobacterium*-mediated transient expression in *N. benthamiana*. Through the incorporation of the suicide exon HyP5SM or the Cre/LoxP recombinase elements, we have enhanced the precision of the inducible system, enabling its utilization in assays highly sensitive to leakiness issues. This design, serving as an alternative to existing chemical-inducible systems, holds the potential for application in other model plants and crop plants.

While DEX- and β-EST-inducible systems stand as the most popular chemical-inducible systems in plant biology, some studies have reported the issue of leakiness associated with these systems (Okuzaki *et al*., 2011; Park *et al*., 2012; Caddell *et al*., 2015; Gonzalez *et al*., 2015). Despite the widespread use of both DEX- and β-EST-inducible systems in many studies, we believe that some attempts utilizing these systems have failed due to leakiness issues and remained unreported. We showed that our improved design of the copper-inducible system can be used to control hypersensitive cell death assays, which is one of the most sensitive assays *in planta*. Therefore, this system has the potential to resolve the leakiness issues of the inducible system on *N. benthamiana* encountered before, opening up the opportunity to successfully implement previously hindered ideas.

Since the first report of using the copper-inducible system *in planta* (Mett *et al*., 1993), it has been adapted for various applications. Nonetheless, the majority of these adaptations proved insufficient due to inadequate induction, discrepancies among plant samples, and issues related to leakiness (Mett *et al*., 1996; Boetti *et al*., 1999; Granger & Cyr, 2000, 2001; Mohamed *et al*., 2001). The copper-inducible system was then improved by Saijo and Nagasawa (2014) and Garcia-Perez et al (2022) through testing combinations of the minimum promoter (5’UTR) and activation domains (Saijo & Nagasawa, 2014; Garcia-Perez *et al*., 2022). Using the existing or newly synthesized MoClo modules, we found that fusing CBS4 promoter to CMV1 or CMV2 5’UTR confers higher reporter gene expression than miniDFR and other 5’UTR modules tested. Furthermore, fusing CUP2 to VP64 or p65 confers higher reporter gene expression compared to other activation domains tested here, although we did not compare it to the VPR (VP64-p65-Rta) tested in Garcia-Perez et al. (2022). While the efficacy of the copper-inducible system and the optimized combinations of trans/cis-elements have to be tested when working on a different plant species, our results together with the results from Garcia-Perez et al. (2022) have convincingly shown that this system is a robust tool to be used in *N. benthamiana* to regulated the expression of trans-genes or endogenous genes.

Given that the copper-inducible system and additional regulatory elements described here were generated following the syntax of MoClo system, laboratories already utilizing MoClo-compatible modules can seamlessly integrate these designs (Weber *et al*., 2011; Engler *et al*., 2014; Gantner *et al*., 2018). Depending on the protein accumulation and tolerance requirements, this system can be employed through various approaches. For experiments where a low level of leakiness is acceptable, directly using the CBS4 promoter with miniDFR or CMV1 5’UTR is possible. In cases where leakiness poses an issue, incorporating mCherry2-HyP5SM as an N-terminal tag module is recommended. While a similar strategy was previously used in Gonzales et al. (2015), our strategy does not require testing the potential insertion sites into the gene of interest. However, it is worth noting that the use of mCherry2-HyP5SM may not be suitable for proteins where N-terminal fusion of tags is undesirable. In such instances, LoxP-mCherry-LoxP or LoxP-mCherry*-LoxP can be employed. While we have demonstrated the system is effective with several NLR/AVR combinations, it is essential to acknowledge that different combinations of NLRs and AVRs may tolerate varying levels of leakiness. Therefore, the optimal combination of modules still needs to be individually tested.

One limitation of this system lies in the fact that plants exhibit a response to copper. While the concentration required to activate the copper-inducible system is lower than the threshold causing visible toxicity effects in *N. benthamiana*, the plants may still exhibit copper responses. Consequently, the system may not be suitable for studying plant responses that involve cross-talk with copper-responsive pathways. Considering copper is commonly used in pesticides, the field application of the inducible system may be incompatible with copper-based fungicides. Nonetheless, the key advantage of the copper-inducible system is also the utilization of Cu^2+^ as the inducer. While potential environmental toxicity exists at high concentrations, it likely has a lesser impact than steroid-based inducers (Kumar *et al*., 2021). Moreover, the production of CuSO_4_ or other types of Cu^2+^ sources is more cost-effective than the production of steroid-based inducers. With a thorough environmental assessment, the system holds the potential for large-scale application in field trials for the production of agriculture-related products (Garcia-Perez *et al*., 2022).

In summary, we improved the existing copper-inducible system by incorporating additional regulatory elements to improve the precision of gene expression. Our study demonstrated applicability in controlling the expression of two commonly used reporter genes, RUBY and luciferase, and conducting cell death assays that are often highly sensitive to leakiness issues. Until now, the application of the copper-based inducible system in plants has been limited to a few studies, and the potential applications, as well as the dynamics of gene expression under this system, remain largely unexplored. Nonetheless, given the promising results we obtained in our analysis, we believe that the copper-inducible system, combined with the additional regulatory elements described here, has the potential for broad application across different plant species to address questions in both fundamental and translational biology.

## Supporting information

Table S1. List of primers used in the study.

Table S2. List of plasmids and sequences used in the study.

## Acknowledgments

We thank Mark Youles (The Sainsbury Laboratory, UK) for providing plasmid modules for molecular cloning. This work was supported by grants from the National Science and Technology Council NSTC-111-2628-B-001-023, NSTC-112-2628-B-001-007, and NSTC-112-2813-C-001-025-B. CHW was funded by the 2030 Cross-Generation Young Scholars Program of the National Science and Technology Council, Taiwan.

## Author contributions

BJC, KYL, YFC, LHC, and CHW designed the research. BJC, KYL, YFC, CYH, FJG, and THS conducted the experiments. BJC, KYL, and CHW analyzed the data. BJC, KYL, and CHW wrote the manuscript. BJC and KYL contributed equally.

## Data availability

The data supporting the findings of this study are available within the article or as supplementary materials. All the materials generated are available upon request.

## Supporting Information

**Figure S1.** The leakiness issue was observed when the transcription factors were expressed in the presence of the matching inducible promoters.

**Figure S2.** Comparisons of different promoters in driving the expression of CUP2-VP16 in the copper-inducible system.

**Figure S3.** Incorporating mCherry2-HyP5SM/OsL5 failed to regulate cell death by inducible R3a and AVR3a properly.

**Table S1.** List of primers used in the study.

**Table S2.** List of plasmids and sequences used in the study.

