## Supplemental Figures S1-3 for "Development of a tightly regulated copper-inducible transient gene expression system in *Nicotiana benthamiana* incorporating suicide exon and Cre recombinase"

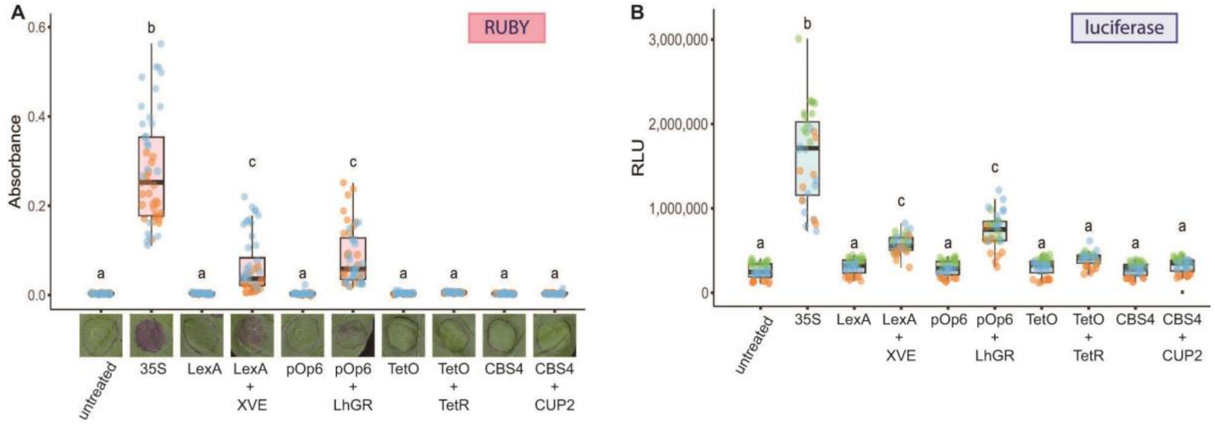


**Figure S1. The leakiness issue was observed when the transcription factors were expressed in the presence of the matching inducible promoters.**

Measurement of RUBY absorbance (A) and luciferase activity (B) controlled by the β-estradiol, dexamethasone, doxycycline, or copper-inducible systems without chemical induction in *N. benthamiana*. RUBY and luciferase reporters were expressed under cauliflower mosaic virus 35S promoter (as positive control), the inducible promoters or the inducible promoters co-expressed with the corresponding transcription factors. For RUBY assays, photographs were taken at 60 hours post-agroinfiltration, followed by RUBY absorbance measurement. Luciferase assays were performed at 48 hours post-agroinfiltration. Dots with different colors represent the results from independent biological replicates. Statistical differences were performed by Tukey’s HSD test ($p<0.05$).

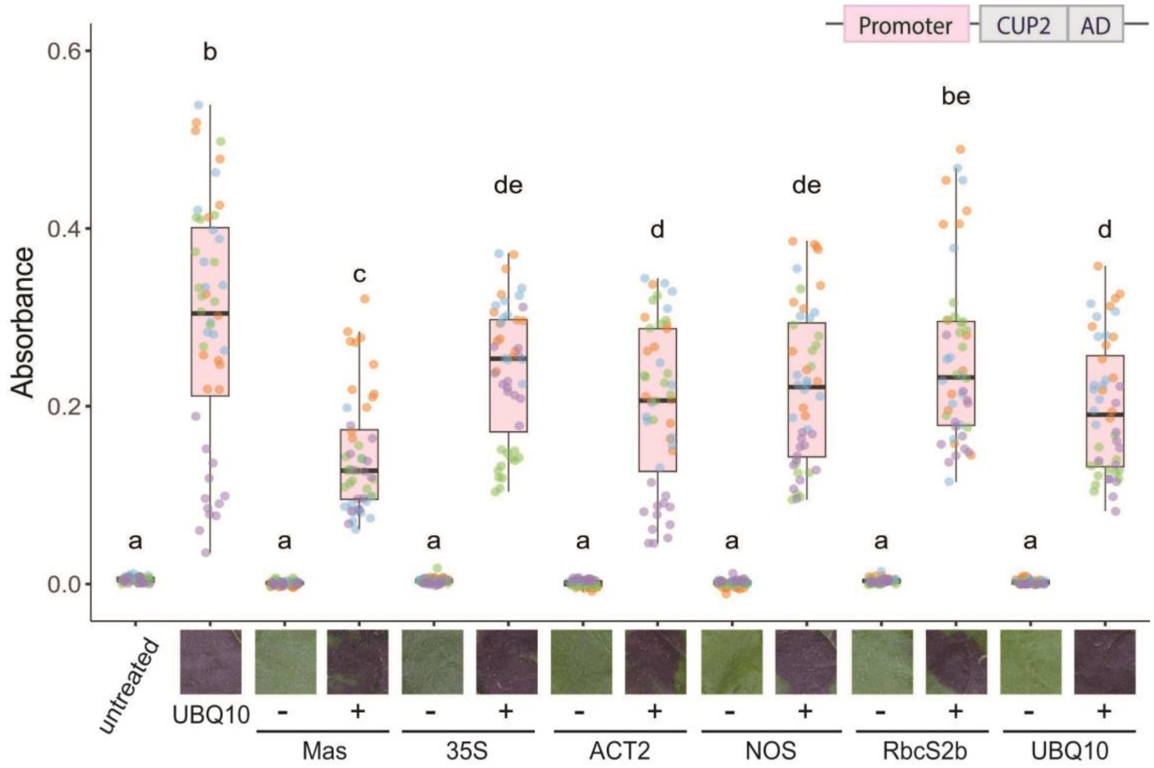


**Figure S2. Comparisons of different promoters in driving the expression of CUP2-VP16 in the copper-inducible system.**

*A. tumefaciens* strains harboring CBS4-miniDFR::RUBY and CUP2-VP16 driven by different promoters were co-infiltrated in four-week-old *N. benthamiana*. The plants were then syringe-infiltrated with 10µM $CuSO_4$ at 24 hours post agroinfiltration. Photographs were taken at 36 hours after $CuSO_4$ infiltration, followed by RUBY absorbance measurement. UBQ10::RUBY was used as a positive control. Dots with different colors represent the results from independent biological replicates. Statistical differences were performed by Tukey's HSD test ($p<0.05$).

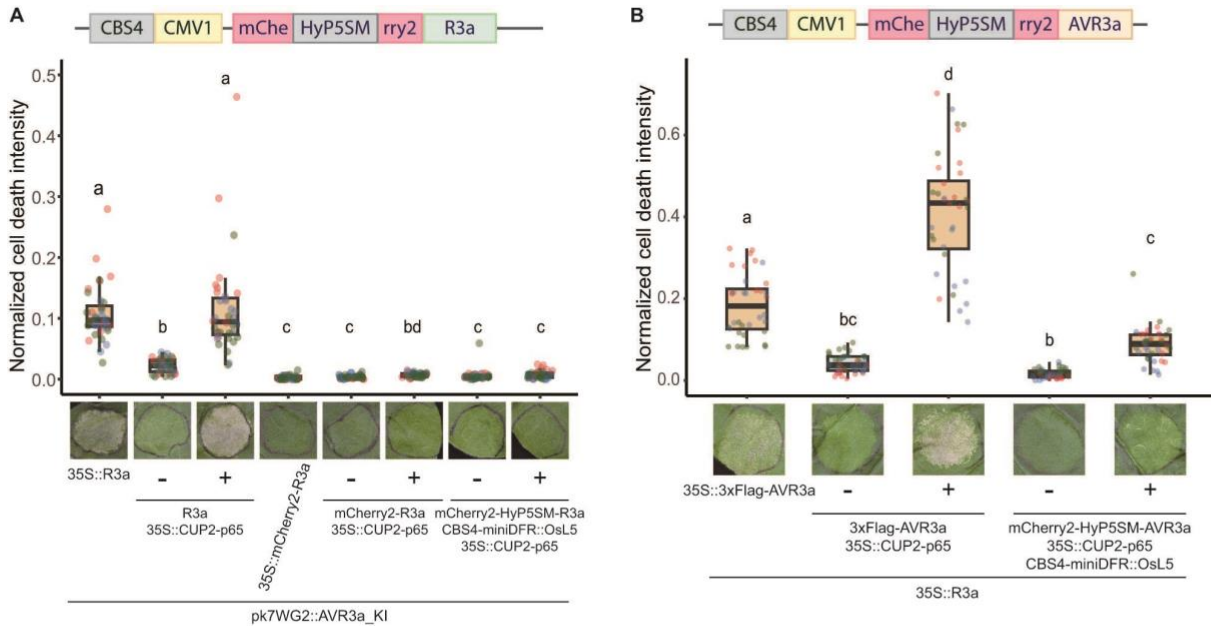


**Figure S3. Incorporating mCherry2-HyP5SM/OsL5 failed to regulate cell death by inducible R3a and AVR3a properly.**

(A) Cell death assay of inducible mCherry2-HyP5SM-R3a/OsL5 with 35S::AVR3a. *A. tumefaciens* strains harboring indicated expression constructs were co-infiltrated in four-week-old *N. benthamiana*. (B) Cell death assay of inducible mCherry2-HyP5SM-AVR3a/OsL5 with 35S::R3a. *A. tumefaciens* strains harboring indicated expression constructs were co-infiltrated in four-week-old *N. benthamiana*. The plants were syringe-infiltrated with 10μM $CuSO_4$ at 24 hours post agroinfiltration. Cell death intensity was quantified using UVP ChemStudio at 36 hours post-copper treatment. For the cell death assays, dots with different colors represent the results from independent biological replicates. Statistical differences were performed by Dunn's test ($p<0.05$).
